# Moisture and material shape microbial communities in the built environment through disturbance–productivity relationships

**DOI:** 10.1101/2025.11.25.690005

**Authors:** Kaelyn Nannini, Dennis D. Krutkin, Scott T. Kelley, Adrian Ortiz-Velez, Jada D. Brown, Shawn M. Ogden, Fernanda Terrazas, Nicholas A. Barber

## Abstract

The built environment houses diverse microbial communities whose diversity and composition differ among building materials and environmental conditions. Ecological theory makes predictions about how productivity and diversity shape communities, and experiments in the built environment provide an opportunity to test these. We manipulated moisture (constant or repeated wet–dry cycling) on three common building materials to test predictions about alpha and beta diversity. The most productive material (oriented strand board) supported the highest bacterial alpha and beta diversity, and these diversity levels were reduced by repeated drying disturbances. Diversity patterns for fungi were more variable, with the highest alpha diversity on low–moderate productivity material (gypsum wallboard). Fungal beta diversity was reduced by disturbance on high-productivity material, but increased on the other materials. These patterns were driven largely by members of Bacillaceae, Sphingomonadaceae, and Aspergillaceae that reached high abundances in some treatments. Differences between bacteria and fungi may be due to the scale-dependence of productivity–diversity relationships. Together, these results indicate that disturbances can interact with building materials, in some cases leading to variation in community composition that makes it difficult to predict the conditions under which microorganisms with potential importance to health and safety will occur.

**Importance:** The built environment—the homes, workplaces, vehicles, and other spaces where people spend most of their time—contains an enormous diversity of microorganisms with significance for human wellbeing, yet we know little about the factors shaping these microbial communities. We found that patterns of wetting and drying that mimic indoor leaks on common building materials affects the diversity of bacteria and fungi growing on the materials. But the identity of these microorganisms differed from one building material to another, and this was especially variable with wetting and drying. This means that common disturbances that lead to microbial growth in homes and offices can make it difficult to predict which microbes, including those that represent health threats to people, will occur in the built environment.

## Introduction

The Built Environment (BE) is the diverse collection of human-made structures in the world, such as houses, restaurants, office buildings, and vehicles. Urbanization and industrialization have caused humans to shift towards an indoor environment, with one study showing Americans spend over 90% of their time indoors (Klepeis *et al*. 2001). These indoor environments—including both growth substrate materials and abiotic conditions—present novel selective pressures and habitats for microbial organisms, shaping the communities of microorganisms that humans encounter (Kelley and Gilbert 2013; Gilbert and Stephens 2018). Variation in community diversity and composition may have important consequences for microbial impacts in the BE, not only on building structure via degradation (Viitanen *et al*. 2010; Ling *et al*. 2014), but also to the overall health and well-being of occupants (Miller and McMullin 2014; Dannemiller *et al*. 2016). Nonetheless, understanding of how BE microbial communities assemble is limited because BEs can be highly variable depending on a factors like material type, moisture, pH, temperature, geographic location, and surface location within the same BE (Viitanen *et al*. 2010; Kelley and Gilbert 2013; Chase *et al*. 2016; Gilbert and Stephens 2018). Moisture and high humidity in particular are key factors in inducing microbial growth (Du, Li and Yu 2021), but moisture and humidity patterns vary widely among and within BEs.

Ecological theory provides a framework for predicting microbial community assembly in the BE (Costello *et al*. 2012; Miller, Svanbäck and Bohannan 2018). Although developed primarily for macroecological communities, theoretical and empirical studies have identified both site productivity and disturbance regime as important community drivers (Grime 1973; Kondoh 2001; Haddad *et al*. 2008; Mouillot *et al*. 2013). Alpha diversity often exhibits a hump-shaped relationship along productivity gradients, particularly when measured at a local scale, so that the highest diversity occurs at moderate productivity (Chase and Leibold 2002). This may be because diversity is resource-limited at low productivity and competition-limited at high productivity. However, frequent disturbances may disrupt this competition limitation, allowing more species to coexist in productive environments (Huston 1999) (Fig. 1A). Alternatively, frequent disturbance could filter out disturbance-intolerant species, reducing diversity regardless of productivity. Productivity and disturbance also influence the multivariate composition of communities, again by acting as either potentially strong deterministic filters that limit community membership to particular taxa, or by reducing deterministic drivers and allowing stochastic processes to drive composition. The former of these outcomes may be more likely under low productivity, and the latter in high productivity environments, where other mechanisms, such as priority effects, could also lead to multiple compositional outcomes when disturbances are limited (Harrison *et al*. 2006; Chase 2010). At the other extreme, the resource-limited conditions of low-productivity environments, and high disturbance frequency, are expected to homogenize communities (Chase 2003; Jiang and Patel 2008) (Fig. 1B).

**Fig. 1.**
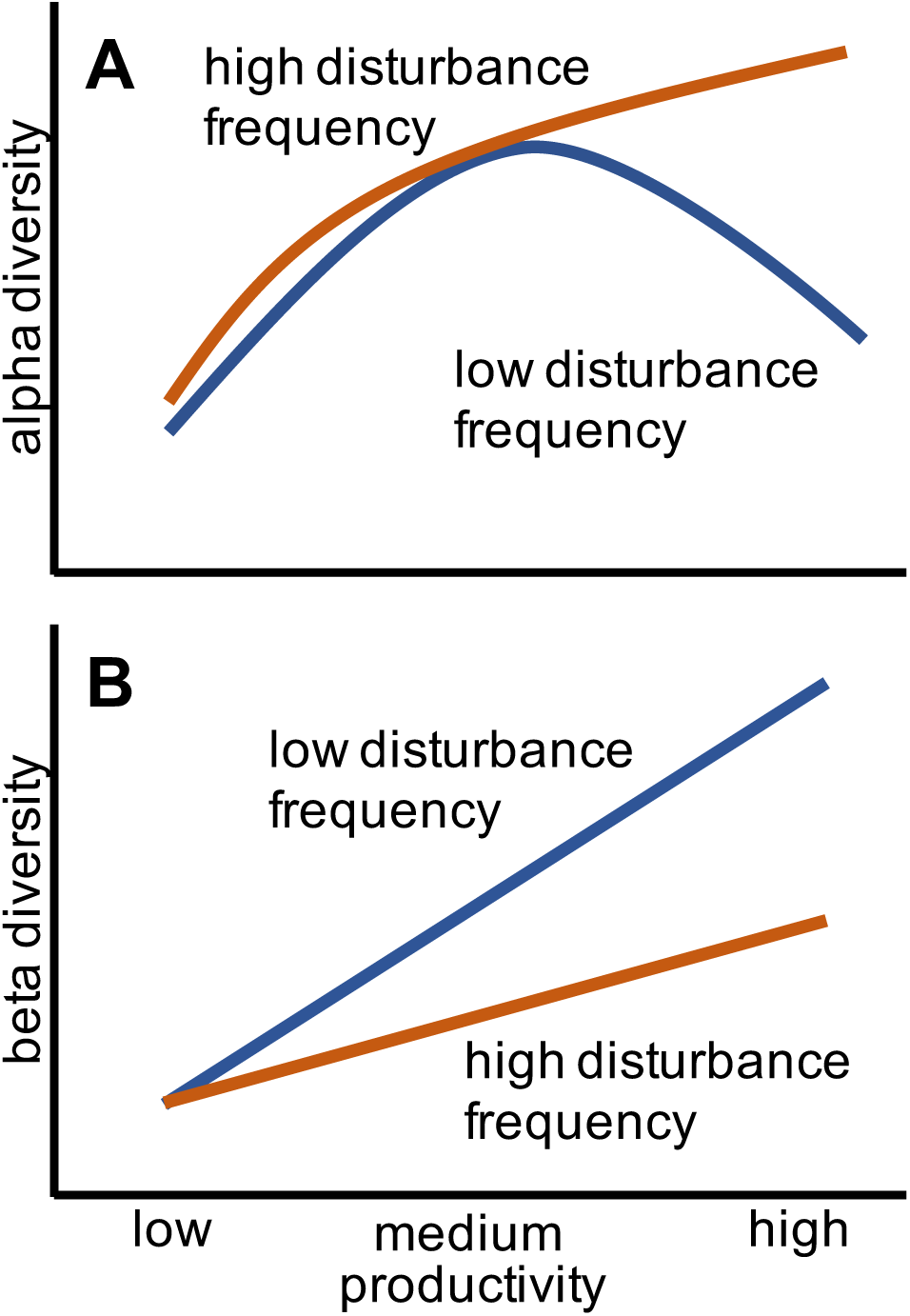
Hypothesized patterns of (A) alpha and (B) beta diversity vs. productivity under high and low frequency disturbance conditions.

In the BE, potential productivity differs among building materials that act as substrates for microorganisms, and drying acts as a disturbance that disrupts microbial communities. Some materials support copious fungal and bacterial growth, and others represent more resource-limited environments in which fewer microbes can establish and grow (Viitanen *et al*. 2010; Lax *et al*. 2019). The most important disturbance regimes in the BE are wetting/drying cycles that occur when building materials become moist from precipitation, plumbing problems, or other causes (Chase *et al*. 2016; Du, Li and Yu 2021). Moisture allows microbes to proliferate following germination of spores and dormant cells (Pasanen *et al*. 2000), turning a dry “microbial wasteland” (Gibbons 2016) into an active community, but subsequent drying represent a stressful filter that limits growth and may influence diversity.

However, our ability to predict how conditions in our built environment shape communities of these microorganisms is limited (Adams *et al*. 2016), in part because much previous research often has focused on observational approaches. When building materials were experimentally maintained at either wet or dry conditions, bacterial and fungal diversity both declined over time, with bacterial diversity lower under consistent moisture (Lax et al. 2019).

But this lower bacterial diversity was likely due to a subset of microbes increasing in abundance and reducing community evenness, compared to dry materials where activity was extremely limited, and the diverse community resulted from dormant microbes that accumulated with dust. In another study that maintained moisture on two different materials, Xu et al. (2020) found differences in bacterial and fungal abundance between them. Notably, this study also demonstrated the importance of incorporating culture-independent counting methods for biomass estimates into community characterization, because some growth patterns were masked or appeared different when examining composition based on relative abundances alone.

In this study, we manipulated disturbance using a wet/dry cycling treatment on three building materials that differ in potential productivity and compared the resulting microbial communities to consistently wet (i.e., low disturbance) communities. We characterized bacterial and fungal communities and used quantitative profiling (combining biomass data and sequence data, *sensu* Vandeputte *et al*. (2017)) to examine effects on alpha and beta diversity. Our design and statistical analysis allowed us to test interactive effects of moisture manipulations and BE materials, which has not been done in similar previous studies. Based on predictions from community assembly theory (Fig. 1), we expected (1) that alpha diversity would increase with productivity under the disturbance treatment but would follow a hump-shaped pattern in low disturbance environments; and (2) that beta diversity, measured as mean pairwise dissimilarity within a treatment combination, would increase with productivity but be reduced by frequent disturbance.

## Methods

### Experimental Treatments and Sampling

Three different building materials were used for this study: medium density fiberboard (MDF), gypsum wallboard (GW), and oriented strand board (OSB). These materials vary in their components and structure and thus represent contrasting environments for microbial growth (i.e., potential productivity). Each material was cut into 5cm x 5cm “coupons.” Materials were UV sterilized, cleaned using 70% ethanol, and allowed to naturally inoculate for a period of 365 days (March 2020-March 2021). Twenty coupons of each material type were positioned in open air on top of a shelf in a laboratory. There was minimal visitation to the laboratory during this time due to the COVID-19 pandemic. Following inoculation, coupons received one of two wetting treatments (n = 10 per treatment per material type). Each constant wet coupon was wetted with 5g of UV-sterilized tap water and placed in separate sealed plastic tub, representing a disturbance-free environment. Coupons undergoing the wet-dry cycling treatment, representing a repeated disturbance regime, were similarly wetted but after one week were drained and allowed to air dry in front of a HEPA filter with the lid resting on top to prevent contamination of debris or particles, then resealed and incubated for a week before sampling and wetting again (Fig. 2).

**Fig. 2.**
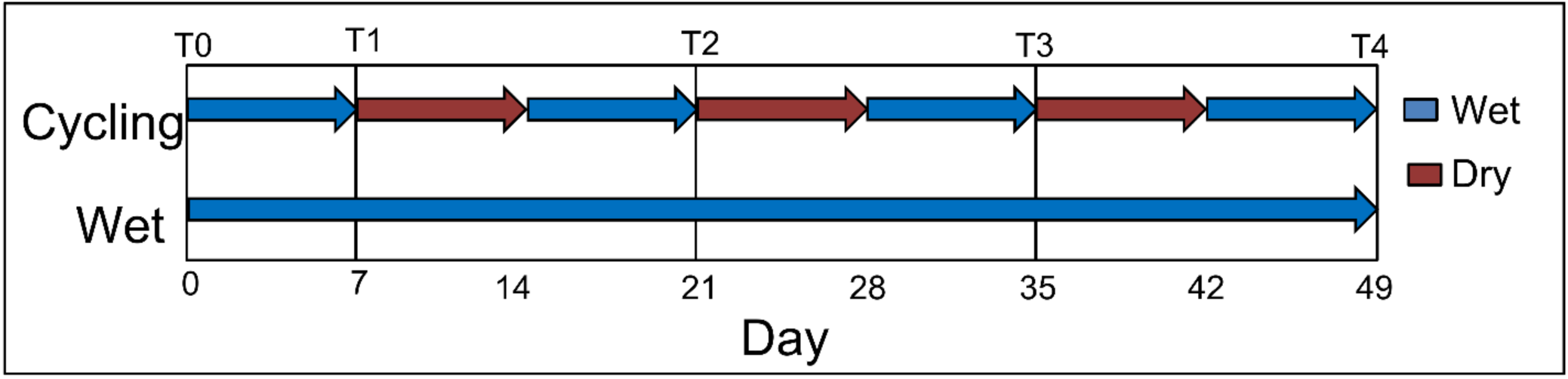
Sampling times for experimental treatments. Wet treatments maintained a consistently moist environment for microbial growth, while cycling treatments alternated through repeated drying period followed by rewetting before each sampling event.

Coupons were sampled at 5 timepoints: week 0 (start of the experiment), week 1, week 3, week 5, and week 7 (T0-T4 respectively), representing three cycles of drying disturbance for cycled coupons. At T0, two coupons of each material/treatment combination were sampled; for the subsequent time points, all coupons were sampled. Samples were taken by opening the container, removing the coupon, and swabbing the surface of the coupon in one direction for approximately 20s. The coupon was then replaced in the tub and sealed. The sample swab was broken off and placed in a 1.5mL microcentrifuge tube containing 700µL of 0.85% NaCl. The microcentrifuge tube was vortexed for 20s. The sample was then aliquoted into separate microcentrifuge tubes: 200uL was aliquoted for microscopy and 150uL was aliquoted for DNA extraction. Samples to be used for microscopy were fixed using a 4% formaldehyde solution and stored at 4°C until further processing.

### Productivity

We used bacterial cell counts, fungal hyphal length, and fungal spore density to estimate microbial productivity. Epifluorescence microscopy was performed for approximately half of the samples at each sampling timepoint. Previously fixed samples were thawed, and 100µL was resuspended in 5mL of 0.02µm filtered water. Samples were vacuum filtered, stained with 1X SYBR Gold, and allowed to incubate for 10 minutes in the dark before washing with 100µL of filtered water for 5 minutes and mounting the filters onto glass slides. For bacterial biomass estimates, the slides were imaged under an Olympus 60X object magnification oil immersion microscope connected to a QImaging Retiga EXi Fast Cooled Mono 12-bit camera. Images were analyzed and counts were estimated using Image Pro software (Media Cybernetics, Rockville, Maryland USA). For fungal counts, a Keyence BZ-X800E Series All-in-One Fluorescence Microscope was used for imaging. Spore counts were estimated using ilastik v1.4.0, and hyphae lengths were analyzed using ImageJ (v2.9.0) (Schneider, Rasband and Eliceiri 2012).

### DNA Extraction, Sequencing, and Processing

DNA extractions were performed for all samples using the QIAGEN DNeasy PowerSoil kit and stored at −80°C until further processing. The extracted DNA was used to amplify target genes in the 16S rRNA gene and ITS region for bacteria and fungi respectively. For all samples, we targeted the V4 region of the bacterial 16S rRNA gene using the 515F and 806R primer pair. The highly variable fungal internal spacer (ITS) region 1 located between the 5.8S and 18S regions was amplified for half the samples using the ITS1 and ITS2 primer pairs. Reverse primers for both sets were barcoded to allow for re-identification of samples after sequencing. PCR conditions for both ITS and 16S amplification consisted of: 1) 94°C for 3min, 2) 94°C for 45s, 3) 50°C for 1min, 4) 72°C for 1:30min 5) Cycling of steps 2-4, 6) 72°C for 10s, 7) 4°C hold. Most samples underwent 30 cycles; however, a small subset of 16S samples underwent 35 cycles due to a failure to low amplification at 30 cycles. PCR was verified via gel electrophoresis on a 1% agarose gel. Sequencing was performed on the Illumina MiSeq platform by the Scripps Research Institute Genomic Center (San Diego, California USA).

Sequencing reads were imported into QIIME2 (11-2022 release) (Bolyen *et al*. 2019) for data processing. Samples were demultiplexed using the q2-cutadapt plugin. The data were denoised using the q2-dada2 plugin to generate sequence variants (SVs). The sequences were trimmed at 250 base pairs for 16S and 240 base pairs for ITS to remove positions with a quality score lower than 30. SVs that were present in less than 4 and 5 samples in 16S and ITS, respectively, were removed from the analysis. The number of samples a feature must be present into to keep was determined mathematical approach for removing zeros developed by (Ortiz-Velez and Kelley 2024). Sequences containing 16S SVs were taxonomically classified using the Silva database v138 (Quast *et al*. 2012) using the classify-sklearn option in the q2-feature-classifier plugin. For sequences containing ITS SVs, a Naïve-Bayes classifier was created using the UNITE fungal database (v9.0) (Abarenkov *et al*. 2024) for taxonomic classification.

### Data Analysis

Bacterial cell counts, fungal hyphae length, and fungal spore density were analyzed using linear mixed models with material type, disturbance treatment, sampling day, and their two- and three-way interactions as categorical fixed factors. Coupon ID was a random factor to account for repeated sampling. Both variables were log-transformed to meet assumptions of residual normality and homoscedasticity. Models were fit using the lmer() function in the lme4 package (Bates *et al*. 2015) of R (R Core Team 2022). Fixed factors were evaluated with type III χ^2^-tests using the Anova() function in the car package (Fox and Weisberg 2019).

We assessed alpha diversity using the Shannon diversity index, which reflects both richness and evenness. Although alpha diversity metrics are usually based on rarefied estimates to account for differences in read counts during sequencing, our read counts varied over several orders of magnitude because the material types we studied vary so widely in productivity (see results, below). Thus, rarefying to the level of samples with minimal read counts is impractical, and choosing an arbitrary minimal read count cutoff would bias sampling by disproportionately discarding certain treatment combinations (particularly disturbed MDF samples, where microbial biomass was very low). Instead, we present unrarefied bacterial and fungal Shannon diversity values. Bacterial and fungal Shannon diversity were analyzed with linear mixed models using the same approach described above for productivity.

We examined beta diversity within treatments by calculating Aitchison’s distance on central log ratio (CLR) transformed read count data, with zeros replaced by a small value (0.01). CLR transformations and Aitchison’s distance are necessary because of the compositional nature of sequence data, which can lead to spurious results when analyzed as simple relative abundances (Gloor *et al*. 2017). We focused on the final time period, after repeated disturbance events had shaped community compositions, and calculated pairwise values for all pairs of coupons within the same material x disturbance treatment combination, for both bacteria and fungi. We analyzed these values using general linear models, with material type, disturbance treatment, and their interaction as fixed factors, using type III F-tests in the Anova() function. We also analyzed bacterial and fungal composition at the final sampling time using perMANOVA with the adonis2 function in vegan, including material, disturbance, and their interaction as predictor variables. We visualized final community composition using NMDS and mean abundance of identified family in material x disturbance treatments and present both relative and quantitative abundances. Quantitative abundances were estimates by multiplying relative abundances by bacterial cell counts and fungal spore density.

### Key Taxa and Interactions

To identify taxa driving differences among materials and treatments, we took two approaches. First, we implemented random forest models using the randomForest package (v4.7.1.1) (Liaw and Wiener 2002). For bacteria, two separate models were created, one for cycling coupons and another for wet coupons to compare the three material types because the combined dataset had a high error rate; fungi were not separated by treatment. Features (SVs) with a Gini index value ≥1.0 were CLR-transformed and used as response variables in linear mixed models with the same parameters used in the biomass analyses.

## Results

### Productivity

Productivity, as measured by bacterial counts, fungal hyphal length, and fungal spore density, differed among materials (Fig. 3A-B, Fig. S1, Table 1). At the end of the experiment, productivity was greatest on OSB and was mostly increased by the wet treatment. Mean bacterial density was about four times greater on OSB than on MDF and GW for constant wet coupons, and six and four times greater, respectively, with the wet cycling disturbance. Disturbance reduced mean bacterial density by 64-77%. Mean fungal length of OSB was over five times greater than MDF and over eleven times greater than GW under the constant treatment. Fungal hyphae and spore patterns were similar but slightly weaker under the disturbed treatment, although MDF had limited fungal productivity with disturbance.

**Fig. 3.**
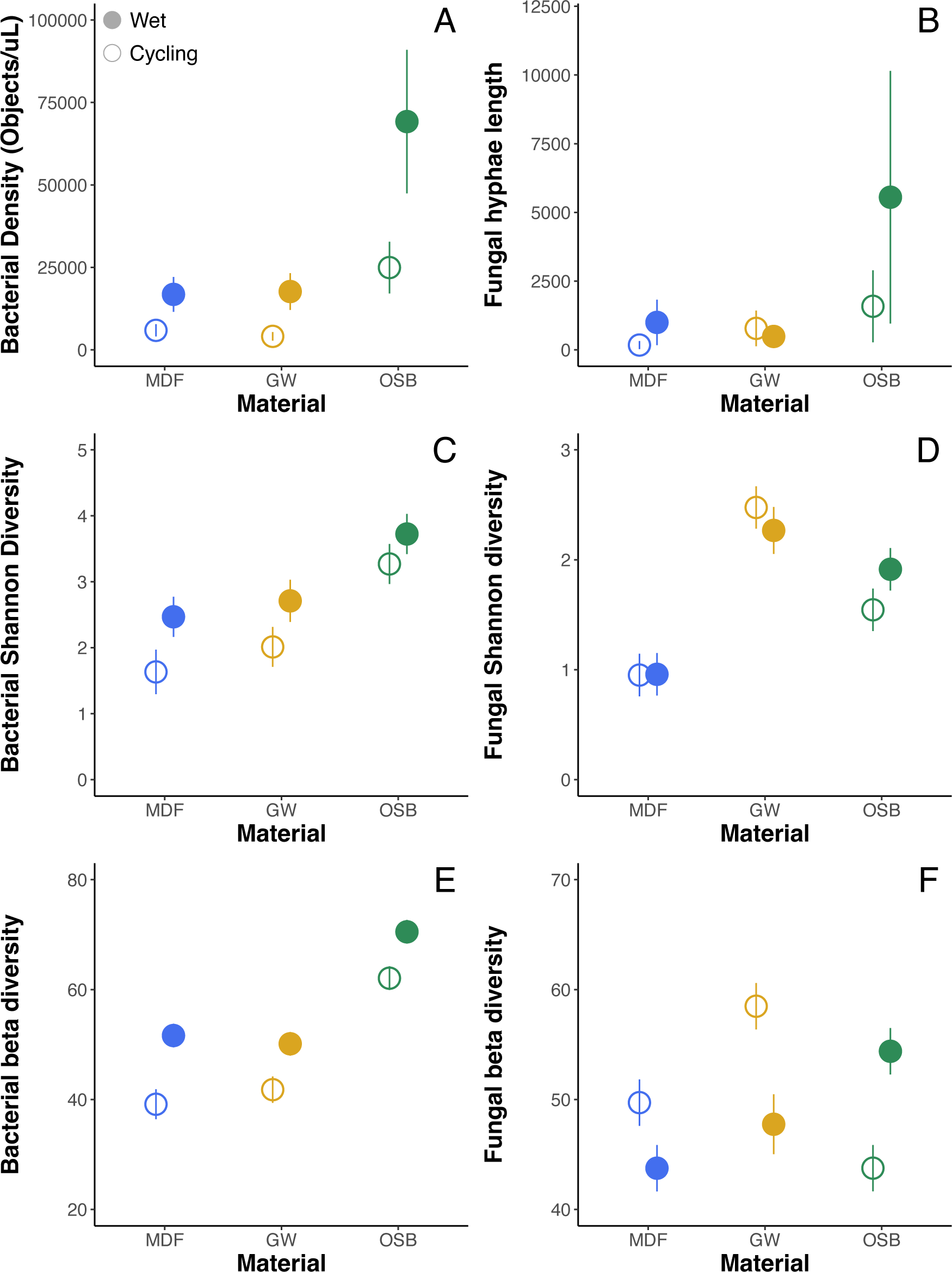
Productivity as measure by (A) bacterial counts and (B) fungal hyphae length on different building materials and under constant (solid symbols) or disturbed (open symbols) wet treatments. (C) Bacterial and (D) fungal Shannon diversity, and pairwise beta diversity (Aitchison’s distance) for (E) bacteria and (F) fungi. Values are estimated marginal means for the last sampling event (± 1 SE).

**Table 1.** Results of linear mixed models analyzing effects of disturbance regime treatment and material type over time on productivity, alpha diversity, and beta diversity.

|  | Bacterial counts |  |  | Fungal hyphae length |  |  |
| --- | --- | --- | --- | --- | --- | --- |
| | $\chi^2$ | df | P | $\chi^2$ | df | P |
| Disturbance | 42.38 | 1 | <0.001 | 1.28 | 1 | 0.258 |
| Day | 78.11 | 3 | <0.001 | 132.13 | 3 | <0.001 |
| Material | 14.83 | 2 | <0.001 | 44.22 | 2 | <0.001 |
| Disturbance x Day | 11.03 | 3 | 0.012 | 1.16 | 3 | 0.763 |
| Disturbance x Material | 1.34 | 2 | 0.512 | 4.14 | 2 | 0.126 |
| Day x Material | 37.08 | 6 | <0.001 | 30.07 | 6 | <0.001 |
| Disturbance x Material x Day | 30.79 | 6 | <0.001 | 9.59 | 6 | 0.143 |

|  | Bacterial Shannon |  |  | Fungal Shannon |  |  |
| --- | --- | --- | --- | --- | --- | --- |
| | $\chi^2$ | df | P | $\chi^2$ | df | P |
| Disturbance | 6.25 | 1 | 0.012 | 0.39 | 1 | 0.535 |
| Day | 9.19 | 3 | 0.027 | 7.42 | 3 | 0.060 |
| Material | 15.82 | 2 | <0.001 | 95.54 | 2 | <0.001 |
| Disturbance x Day | 3.07 | 3 | 0.381 | 5.85 | 3 | 0.119 |
| Disturbance x Material | 3.01 | 2 | 0.222 | 0.07 | 2 | 0.966 |
| Day x Material | 39.15 | 6 | <0.001 | 32.19 | 6 | <0.001 |
| Disturbance x Material x Day | 14.06 | 6 | 0.029 | 14.72 | 6 | 0.023 |

|  | Bacterial Beta diversity |  |  | Fungal Beta diversity |  |  |
| --- | --- | --- | --- | --- | --- | --- |
|  | F | df | P | F | df | P |
| Disturbance | 26.96 | 1 | <0.001 | 9.74 | 1 | 0.003 |
| Material | 57.04 | 2 | <0.001 | 12.37 | 2 | <0.001 |
| Disturbance x Material | 0.50 | 2 | 0.609 | 13.01 | 2 | <0.001 |

### Alpha diversity

Bacterial and fungal alpha (Shannon) diversity differed over time among both disturbance regimes and materials, as indicated by a significant three-way interaction (Table 1). Bacterial diversity at the end of the experiment was greatest on OSB and reduced by the disturbance treatment (Fig. 3C). Thus, bacterial diversity increased with productivity under disturbance, as predicted. However, our hypothesis that high-productivity samples would have low alpha diversity in the absence of disturbance was not supported. Fungal alpha diversity showed different patterns, with the highest Shannon diversity on GW sampled with disturbance (Fig. 3D), and OSB diversity intermediate between the other two material types.

### Beta diversity

Beta diversity, as measured by mean pairwise Aitchison distance at the end of the experiment, was significantly affected by both material type and disturbance treatment (Table 1). For bacterial communities, beta diversity was greatest on OSB and reduced by the wet cycling disturbance treatment (Fig. 3E). However, there was no interaction between these treatments, as hypothesized, so the effect of cycling was similar among the three material types. Fungal beta diversity patterns were different, with disturbance increasing beta diversity on both MDF and GW but reducing it on OSB (Fig. 3F). perMANOVA showed that taxonomic composition of bacterial communities was significantly shaped by material, disturbance, and their interaction, while fungal communities only differed significantly by material (Table 2, Fig. 4). Family composition of bacterial and fungal communities differed by material type and disturbance treatment (Fig. 5), with wet and cycling treatments different in composition most on GW, although there were also differences in the relative abundances of fungal families between treatments on OSB.

**Fig. 4.**
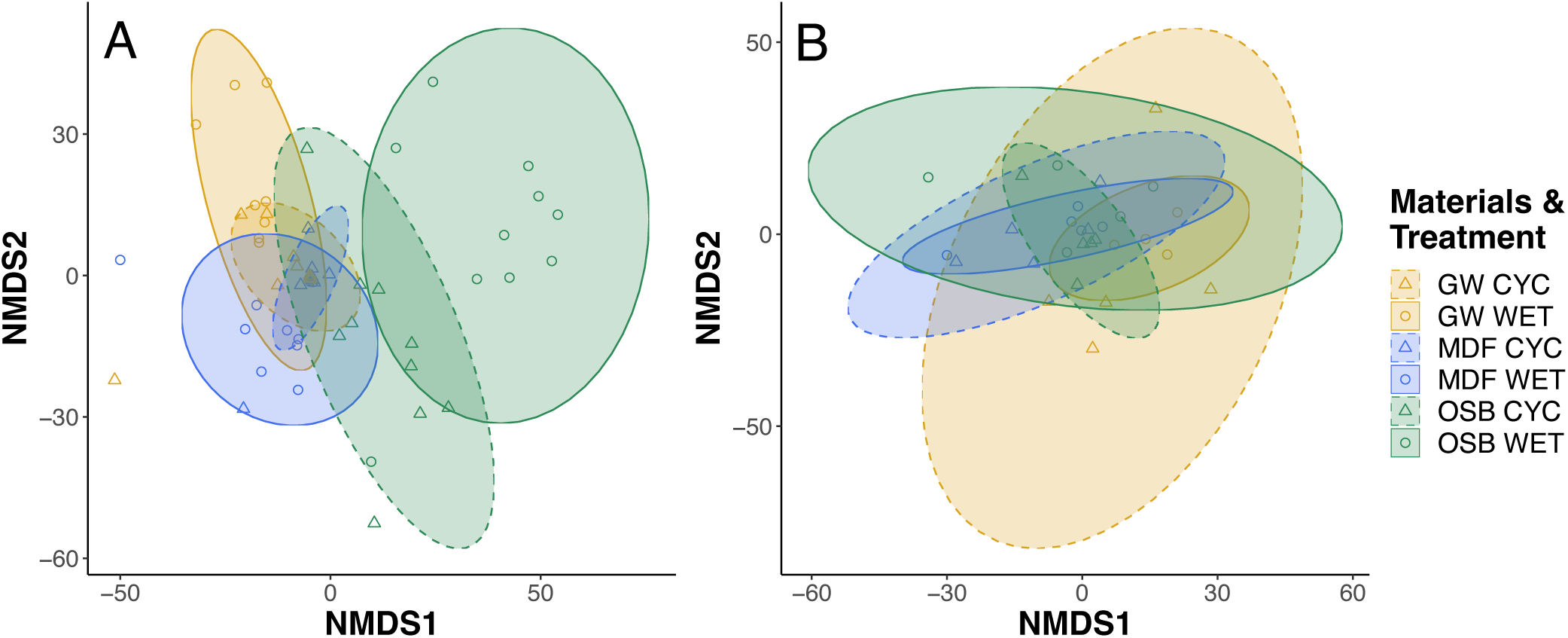
NMDS plots of (A) bacterial and (B) fungal communities at end of experiment. Solid line ellipses and circles represent constant wet treatment, and dashed line and triangles represent cycling wet treatment.

**Fig. 5.**
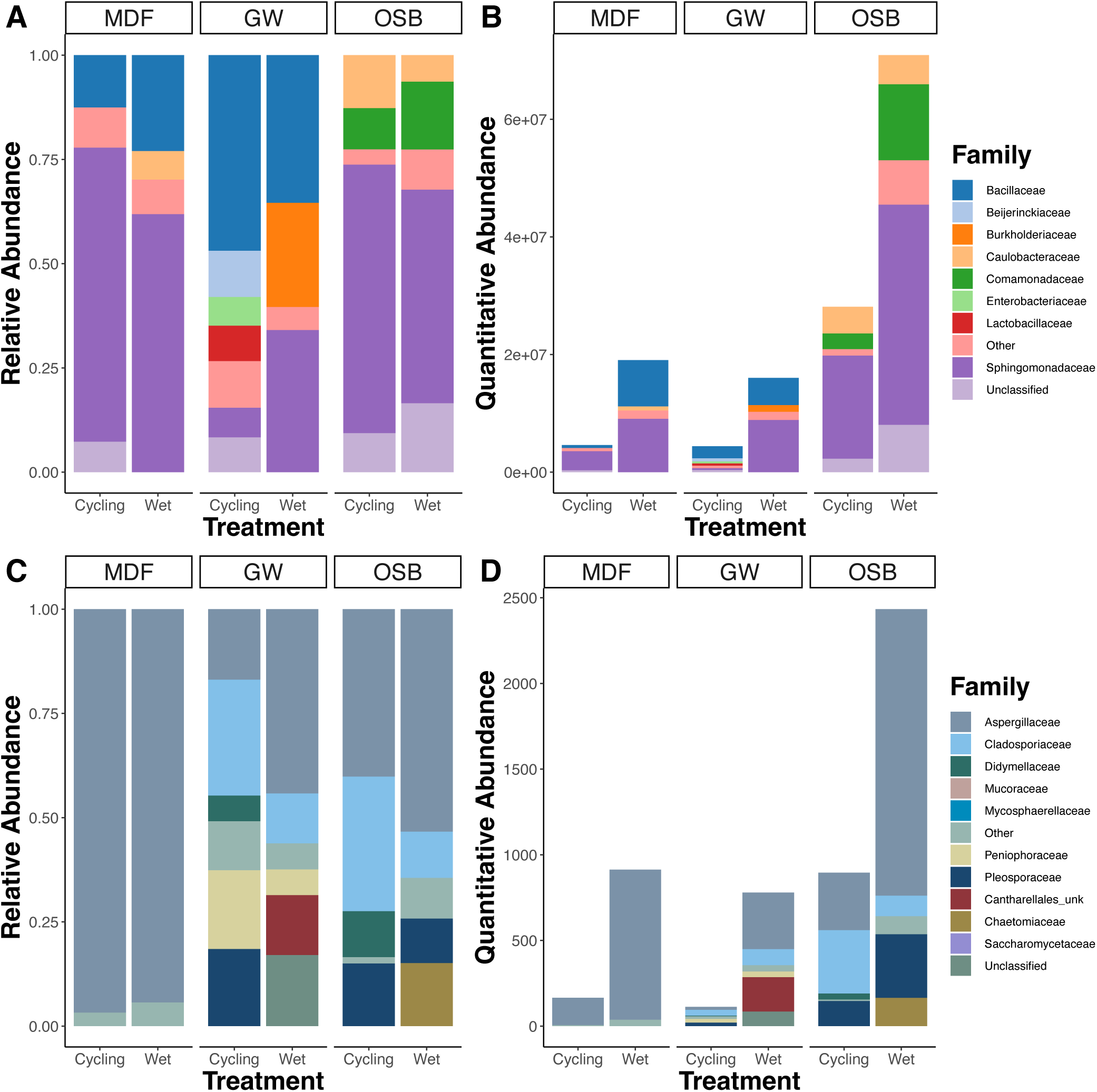
Family composition of (A, B) bacterial and (C, D) fungal communities on building materials by disturbance treatment at end of experiment.

**Table 2.** Results of perMANOVA on bacterial and fungal community composition at the end of experiment by material type and disturbance treatment.

|  | Bacterial composition |  |  |  | Fungal composition |  |  |  |
| --- | --- | --- | --- | --- | --- | --- | --- | --- |
|  | F | df | P | R<sup>2</sup> | F | df | P | R<sup>2</sup> |
| Disturbance | 3.99 | 1 | <0.001 | 0.05 | 1.13 | 1 | 0.210 | 0.04 |
| Material | 9.02 | 2 | <0.001 | 0.23 | 1.57 | 2 | 0.001 | 0.11 |
| Disturbance x Material | 3.39 | 2 | <0.001 | 0.09 | 0.89 | 2 | 0.815 | 0.06 |
| Residual |  |  |  | 0.64 |  |  |  | 0.79 |

### Key Taxa

The two bacterial random forest models identified a combined 13 SVs with a Gini index over 1.0, 9 *Bacillus* spp. and 4 *Sphingomonas* spp. Linear mixed effect models using each of these SVs as a response variable showed significant differences in abundance among material types (Disturbance x Material x Day interaction, *P* < 0.05 for all but two bacterial SVs), with *Bacillus* SVs consistently more abundant on GW and *Sphingomonas* SVs generally reaching higher abundances on OSB (Fig. 6). The fungal random forest model identified 6 SVs: 3 *Penicillium* spp., 2 *Cladosporium* spp., and 1 *Aspergillus* sp. Abundances of these SVs were significantly different among material types (all P < 0.02), with two *Penicillium* SVs (*P. brevicompactum* and *P. sumatraense*) and the *Aspergillus* SV generally more common on MDF. In contrast, two *Cladosporium tenuissimum* SVs were less abundant on MDF.

**Fig. 6.**
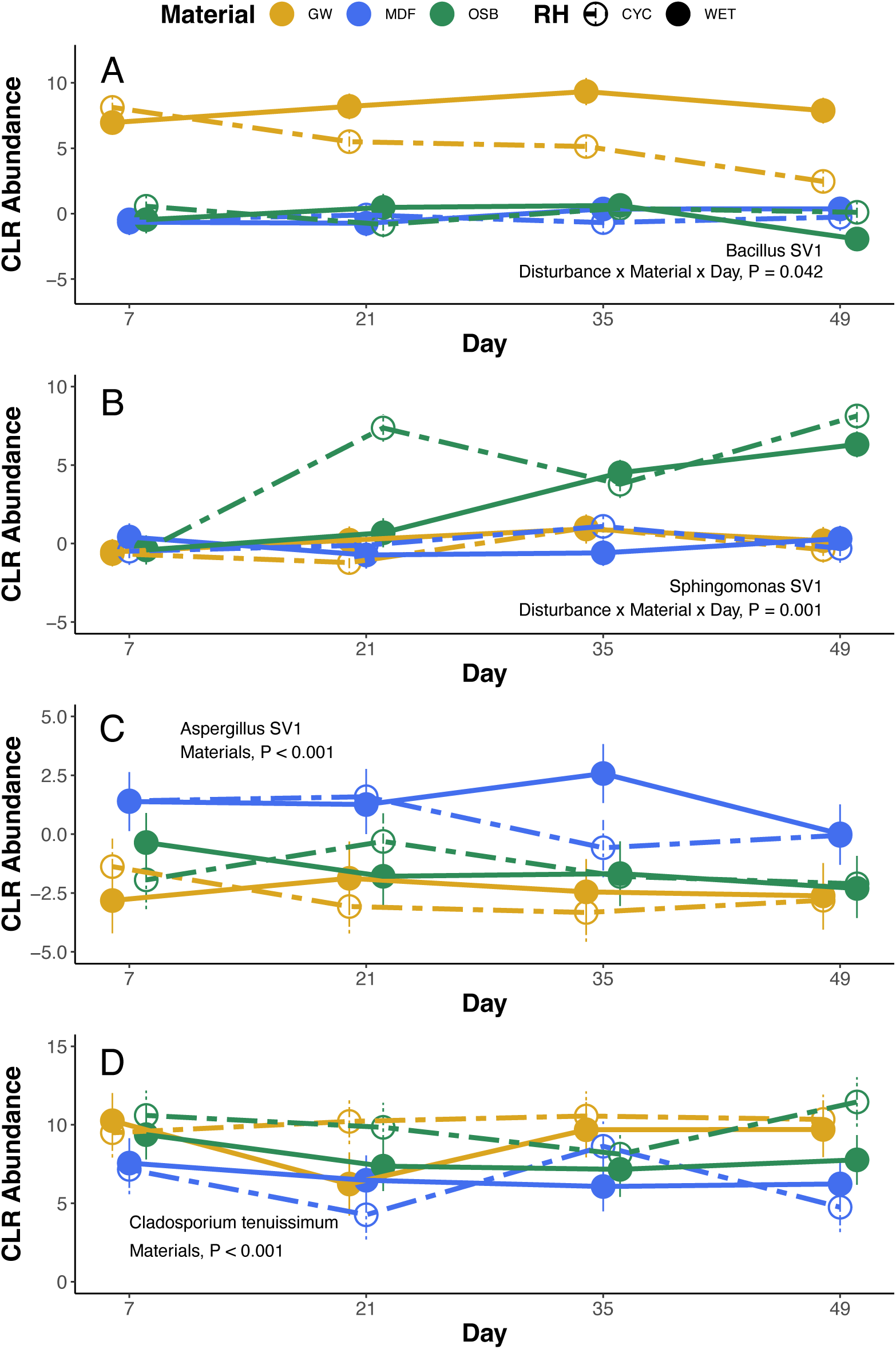
Abundances of select focal SVs identified by random forest analysis as distinguishing material types. (A) *Bacillus* SV1 and (B) *Sphingomonas* SV1 are typical of the other *Bacillus* and *Sphingomonas* SVs, respectively. (C) *Aspergillus* SV1 and (D) *Cladosporium tenuissimum* illustrate differences in abundance on OSB. Abundances are CLR transformed estimated marginal means (± 1 SE).

## Discussion

We examined bacterial and fungal communities on three common building materials under either constant moisture conditions or wet–dry disturbance cycles. Bacterial and fungal growth were both significantly higher on one building material, OSB, indicating it provides high potential productivity as a substrate for microbial communities (Fig. 3A-B). Although we predicted that alpha diversity would follow a unimodal relationship in the absence of disturbance and increase under repeated disturbances, bacterial and fungal diversity patterns differed.

Bacterial alpha diversity responded similarly to disturbances at all productivity levels, maintaining high diversity on OSB, while fungal alpha diversity was highest on GW, a low- to moderate-productivity substrate. Our expectation that a positive relationship between beta diversity and productivity would be strongest under low disturbance was partially supported: bacterial beta diversity was greatest on OSB, but beta diversity was similarly reduced by disturbance on all materials. Fungal beta diversity was increased by disturbance on low-productivity materials but reduced on OSB. Below, we discuss possible explanations for these patterns, which microbial taxa may have driven the results, and the implications for understanding and predicting microbial communities the BE.

### Alpha and beta diversity

A unimodal relationship between productivity and diversity is often assumed because more species are supported along increasing resource gradients, but at the highest productivity levels, a competitively dominant subset of the community can limit diversity (Huston 1999). We expected that disturbances, in the form of repeated drying, would limit this competitive exclusion, permitting higher diversity on the most productive material. However, this did not occur, and material–disturbance interactions on Shannon diversity were fairly weak, especially for bacteria (Fig. 3C-D). Disturbance generally reduced alpha diversity, including for both bacteria and fungi on OSB, the most productive material. One possible explanation for these alpha diversity patterns is the scale-dependence of productivity–diversity relationships.

Unimodal patterns may emerge when assessed at local scales, while at larger scales diversity increases consistently with productivity: Chase and Leibold (2002) found a hump-shaped relationship when sampling individual ponds within watersheds, but a linear relationship when comparing among entire watersheds. Similarly, especially given the size differences between single-celled bacteria and much larger multicellular fungi, we may have been sampling bacteria at a regional scale (essentially pooling across many local environments with each swab across a coupon), whereas fungal samples better represented the local coupon scale at which interspecific interactions occur. Characterizing bacterial communities at the local scale may require much more localized sampling within coupons, although it may be challenging to sequence such low-biomass samples. An additional caveat is that, even if unimodal productivity–diversity relationships exist, it can be difficult for investigators to know where along the productivity gradient the studied environments lie. That is, OSB was clearly the most productive of these three materials, but it could still be quite low relative to other growth surfaces containing more (or more diverse) substrates in the BE or elsewhere (Gibbons 2016).

We also expected variation in community composition among coupons, or beta diversity, to depend on material type and disturbance treatment. Both low productivity and frequent disturbance can function as environmental filters, limiting community membership to species with tolerance for limited resources or resilience against disturbances (Chase 2003; Jiang and Patel 2008). These processes would result in a consistent, limited set of species persisting in these environments and low among-community beta diversity. The prediction that beta diversity would increase with productivity was partially supported: bacterial beta diversity was highest on high-productivity OSB regardless of disturbance, and fungal beta diversity was highest on OSB when disturbance was absent (Fig. 3E-F). These effects are visible in the wide dispersion of OSB samples in ordinations (Fig. 4A-B). However, under frequent disturbances, fungi on OSB had surprisingly limited compositional dispersion (Fig. 3F & 4B), indicating that a limited number of taxa may have been able to take advantage of the resource availability following disturbances and reach high abundances on these coupons. The identity of these taxa could have important practical implications if they tend to be species that cause physical degradation of their substrate or cause health problems.

### Taxonomic composition

Understanding which taxa drove the diversity patterns discussed above may help further reveal the ecological mechanisms shaping these microbial communities. Importantly, we show how combining relative abundances of bacteria and fungi with counts of cell (bacteria) or spore (fungi) density to generate quantitative abundances, gives a clearer picture of community differences (Xu *et al*. 2018).

Within bacterial communities, several *Sphingomonas* SVs were typical of OSB in general, and members of Sphingomonadaceae more broadly comprised much of the bacterial communities, except on GW subject to the drying disturbance. If *Sphingomonas* are particularly susceptible to drying, then the thin surface paper of GW may have limited their persistence in comparison to wet samples or the other two wood-based materials that may maintain moisture longer. Bacillaceae were also significant components of bacterial communities, except on OSB. Even under drying disturbances, they persisted on MDF and GW coupons, albeit at low-biomass. Several *Bacillus* SVs were also identified as distinguishing GW from the other materials. In a similar experiment, Lax et al. (2019) found that *Bacillus* reached high relative abundances on consistently wet GW but remained rare on other materials or when coupons were kept dry. The persistence of *Bacillaceae* on wet-cycling GW and MDF coupons in our experiment suggests that they may be tolerant of low-productivity and high-disturbance conditions, possible due to their well-known ability to form spores.

In fungal communities, two *Penicillium* SVs and an *Aspergillus* SV were identified as characteristic of MDF coupons. These and other Aspergillaceae were the dominant components of MDF communities, although they were a large part of GW and OSB communities, reaching especially high abundances in wet conditions. These findings align with other studies of the BE, in which Aspergillaceae is often common or dominant. On MDF and GW, the greater relative and quantitative abundance of Aspergillaceae on wet coupons was associated with reduced beta diversity among these communities, whereas the higher quantitative abundance due to wet treatment on OSB increased beta diversity. One explanation is that the same SVs reached high abundances on the low-productivity MDF and GW coupons, homogenizing these communities, while on OSB different taxa might have become common and driven the high inter-site compositional variation. If mechanisms like priority effects or ecological drift (Chase 2007; Fukami 2015) are more important under higher productivity, that could have driven the opposite disturbance effects on OSB. This pattern might be widespread on other building materials that, like OSB, are prone to copious microbial growth.

One important caveat to these interpretations is that the idea of environmental filtering as a structuring force in communities may overlook biotic interactions (Kraft *et al*. 2015). The network analysis of Lax et al. (2019) found potential interactions between bacteria and fungi, and the production of antibiotics by some fungi could promote resistant bacteria through apparent competition. Thus, dominance by an antibiotic-producing fungus could represent an additional disturbance on entire bacterial communities but a positive feedback for resistant bacteria, especially given the potential scale differences of bacterial and fungal communities discussed above.

### Implications

Microbial growth in the BE can represent an economic, safety, and health threat because of the potential for material degradation, toxin production, and pathogen spread (Viitanen *et al*. 2010; Miller and McMullin 2014; Gilbert and Stephens 2018). Shifts in the diversity and composition of microbial communities could promote or suppress the agents of these negative impacts, but the community-level interactions that determine this are poorly known. Changing precipitation patterns under climate change may alter wetting frequencies or patterns, driving the sorts of community variations we document in this study. For example, consistent moist conditions, which could arise in areas where rainy seasons last longer or rainfall events become more frequent, promote many *Aspergillus* spp., including those that produce toxins that threaten human health (Prenafeta-Boldú, Medina-Armijo and Isola 2022). Further, we have shown here that different wetting regimes may interact with BE materials to constrain (reduced beta diversity) or amplify (increased beta diversity) compositional variation. This was evident in fungal communities on GW, which had higher beta diversity under the wet cycling treatment. High beta diversity, or wide variation in community structure among samples experiencing the same conditions, means that it may be very difficult to predict important aspects of community composition—such as the presence of potentially threatening microbes—under the sort of wetting and drying cycles that can occur with roof leaks and periodic precipitation.

Microbial-resistant building materials, such as “mold-free” GW or material coatings that limit bacterial and fungal growth, hold promise for limiting potential BE degradation or health effects (Wu and Wong 2020). But the same issue of low predictability of which particular taxa succeed on these materials applies: although we found that lower productivity of bacterial communities was associated with lower alpha and beta diversity (i.e., more constrained and consistent combinations of bacteria taxa), the high beta diversity of fungal communities on low-growth GW means that even when fungal growth is limited, the species that are able to succeed may be vary from one sample to another, even if they were initially inoculated in the same location. Studies targeting the occurrence of particularly harmful fungi (or bacteria) may help reveal if there are specific conditions, including broader community context, that require careful attention to reduce threats in homes, schools, workplaces, and other parts of the BE.

## Acknowledgments

This work was supported by SDSU University Grant Program and the SDSU College of Sciences.

---

**Fig. S1.**
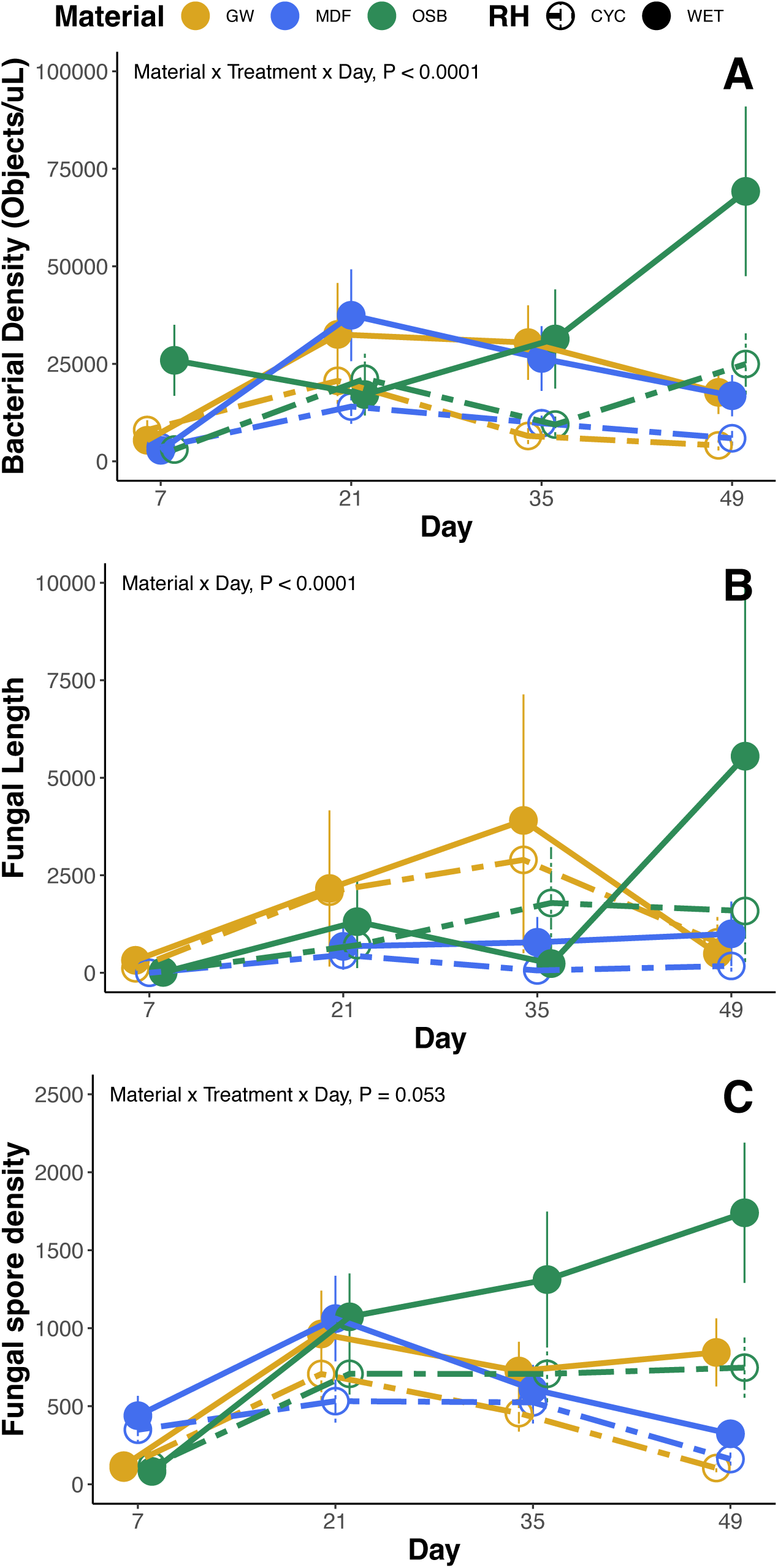
Mean (A) bacterial density, (B) fungal hyphae length, and (C) fungal spore density by material and disturbance treatment over time.

**Fig. S2.**
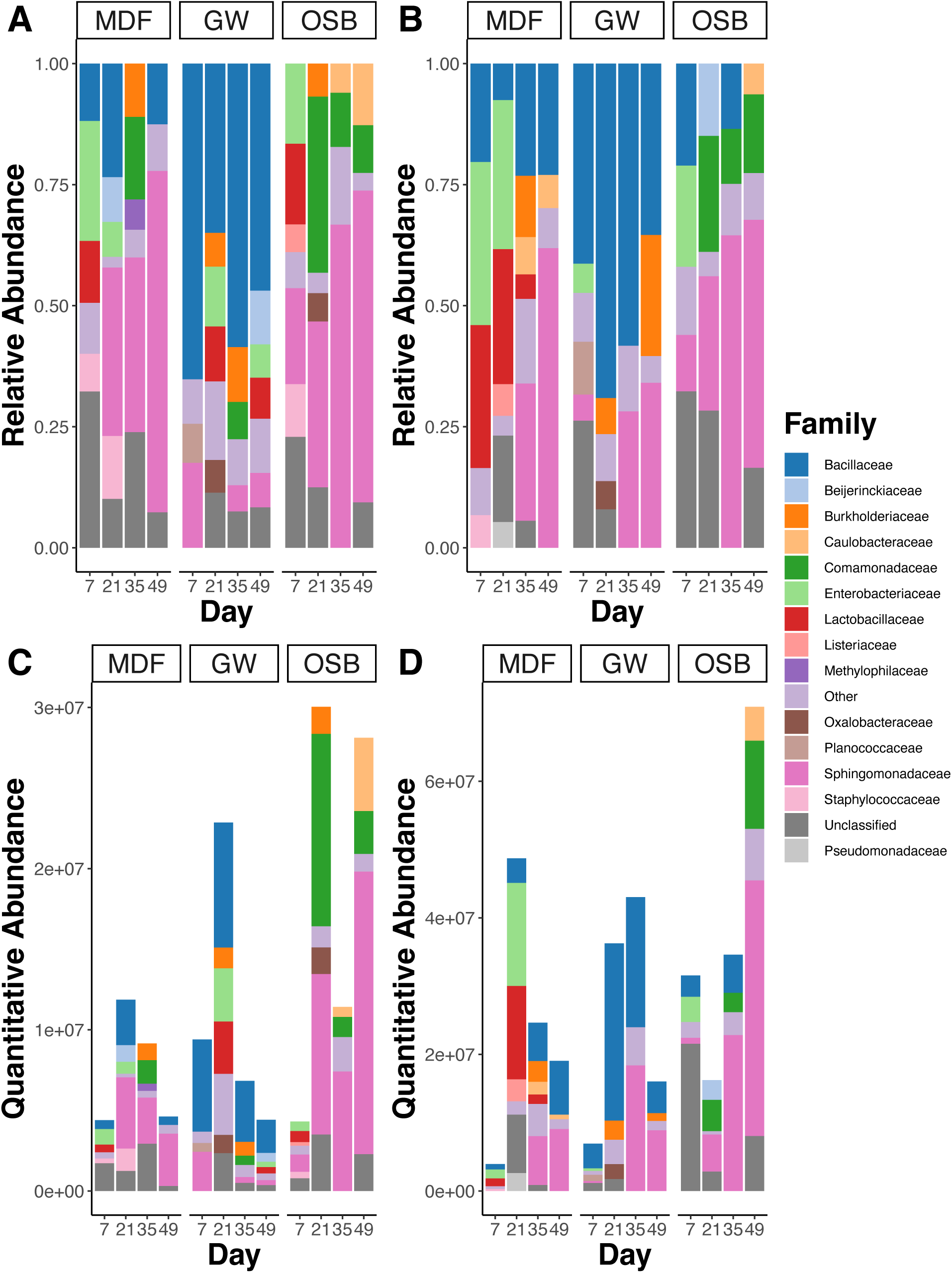
Family composition of bacterial communities on building materials over time, separated by (A, C) constant or (B, D) disturbed wet treatments. Note the different scales in C and D.

**Fig. S3.**
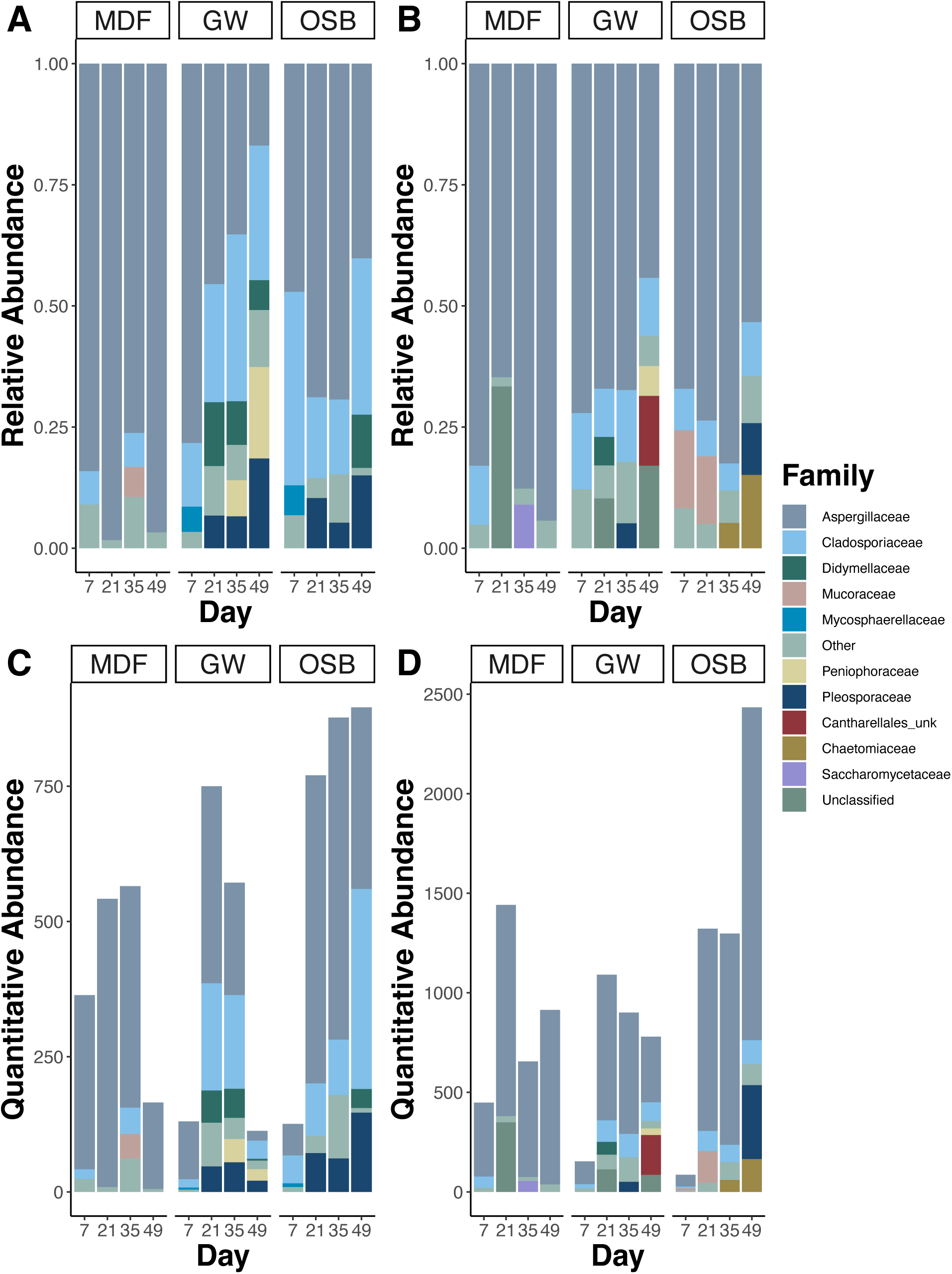
Family composition of fungal communities on building materials over time, separated by (A, C) constant or (B, D) disturbed wet treatments. Note the different scales in C and D.

